# A few stickleback suffice for the transport of alleles to new lakes

**DOI:** 10.1101/713040

**Authors:** Jared Galloway, William A. Cresko, Peter Ralph

## Abstract

Threespine stickleback populations provide a striking example of local adaptation to divergent habitats in populations that are connected by recurrent gene flow. These small fish occur in marine and freshwater habitats throughout the Northern Hemisphere, and in numerous cases the smaller freshwater populations have been established “de novo” from marine colonists. Independently evolved freshwater populations show similar phenotypes that have been shown to derive largely from the same standing genetic variants. Geographic isolation prevents direct migration between the freshwater populations, strongly suggesting that these shared locally adaptive alleles are transported through the marine population. However it is still largely unknown how gene flow, recombination, and selection jointly impact the standing variation that might fuel this adaptation. Here we use individual-based, spatially explicit simulations to determine the levels of gene flow that best match observed patterns of allele sharing among habitats in stickleback. We aim to better understand how gene flow and local adaptation in large metapopulations determine the speed of adaptation and re-use of standing genetic variation. In our simulations we find that repeated adaptation uses a shared set of alleles that are maintained at low frequency by migration-selection balance in oceanic populations. This process occurs over a realistic range of intermediate levels of gene flow that match previous empirical population genomic studies in stickleback. Examining these simulations more deeply reveals how lower levels of gene flow leads to slow, independent adaptation to different habitats, whereas higher levels gene flow leads to significant mutation load – but an increased probability of successful population genomic scans for locally adapted alleles. Surprisingly, we find that the genealogical origins of most freshwater adapted alleles can be traced back to the original generation of marine individuals that colonized the lakes, as opposed to subsequent migrants. These simulations provide deeper context for existing studies of stickleback evolutionary genomics, and guidance for future empirical studies in this model. More broadly, our results support existing theory of local adaptation but extend it by more completely documenting the genealogical history of adaptive alleles in a metapopulation.

## Introduction

The canonical model for the genetics of adaptation has long been the sequential fixation of new mutations [Smith and Haigh, 1974, Endler, 1977, Orr, 2005]. While it has proved to be a useful baseline for understanding the genetic variation we see in species today, this model is now rightfully understood as incomplete for many species in nature that have more complicated population structures [Lai et al., 2019, Schrider and Kern, 2017]. In particular, empirical studies have increasingly identified the need to more deeply incorporate standing genetic variation into adaptation dynamics for metapopulations inhabiting an array of habitats [Hermisson and Pennings, 2005, Barrett and Schluter, 2008]. Populations experiencing diverse selective pressures while still exhibiting significant gene flow often result in more complex genomic signals that are still not fully understood [Charlesworth, 1997, Charlesworth et al., 2003, Nosil et al., 2009, Flaxman, 2013, Samuk et al., 2017]. Concurrently, a growing number of empirical studies have identified instances of convergent evolution using standing genetic variation [Schrider and Kern, 2017, Barrett and Schluter, 2008, Nelson and Cresko, 2018, Nelson et al., 2019, Bassham et al., 2018]. However, it is still not clear how variation in evolutionary processes – such as gene flow, recombination, selection and mutation – can promote the maintenance and re-use of standing genetic variation, particularly during colonization and adaptation to new environments [Nelson and Cresko, 2018, Pritchard, 2010, Yeaman and Whitlock, 2011, Schrider and Kern, 2017]. It is similarly unclear whether variation in these evolutionary processes can determine the genetic architecture of evolving traits via standing genetic variation. Theoretical work is mostly limited to one or two loci, or relies on approximations of uncertain validity [e.g., Slatkin, 1975, Petry, 1983, Barton and Bengtsson, 1986, Yeaman and Whitlock, 2011, Ralph and Coop, 2015]. Therefore, detailed simulation models of specific systems can provide an important complement to empirical studies in the lab and field, by helping us understand precisely how standing genetic variation might fuel local adaptation, and what genomic signals we can expect to see.

The ancestral marine form of threespine stickleback fish has given rise to millions of independently derived freshwater populations in recently de-glaciated regions around the Northern Hemisphere [Hunt et al., 2008, Thompson, 1997, Cresko et al., 2007]. This model organism has provided some of the earliest data showing the heterogeneous nature of divergence across genomes and the much more extensive use of standing genetic variation than once thought [Schluter and Conte, 2009, Roesti et al., 2014, Nelson and Cresko, 2018, Bassham et al., 2018, Nelson et al., 2019, Hohenlohe et al., 2010, Terekhanova et al., 2014, Marques et al., 2016]. While geographic isolation often prevents direct migration between freshwater populations, stickleback in them frequently evolve similar phenotypes [Cresko et al., 2004, Colosimo et al., 2004]. The most recent evolutionary genomic studies on stickleback document that while the overall dynamic of local adaptation to marine and freshwater habitats has been occurring for millions of years [Nelson and Cresko, 2018], independent local adaptation of marine individuals to freshwater environments has been observed to take place in just tens of generations [Terekhanova et al., 2014, Lescak et al., 2015, Bassham et al., 2018]. For example, in 1964 the Great Alaskan Earthquake caused an uplift of many islands and coastal regions throughout the Gulf of Alaska. Studies of stickleback populations on uplifted Middleton Island showed that newly created freshwater ponds were invaded by the surrounding marine population of stickleback which evolved the freshwater syndrome of phenotypes in less than 50 years [Lescak et al., 2015, Bassham et al., 2018]. Amazingly, the portions of the genomes of these populations that showed increased divergence from the oceanic population mirrored those previously found to differ between ocean and freshwater populations that that have been geographically separated for thousands of years [Hohenlohe et al., 2010, Bassham et al., 2018].

But how can evolution occur at such a rapid pace? Waiting for new mutations to arise in each lake or pond would take much longer than the decades since the Alaskan earthquake with any plausible target size. Even more improbable is having divergence cluster in such similar genomic regions across independent populations. An alternative hypothesis is that the majority of alleles important for freshwater adaptation are maintained in the marine individuals due to recurrent gene flow from freshwater back in to marine populations. Schluter and Conte [2009] proposed a conceptual model they termed the “transporter” hypothesis to describe the process by which alleles beneficial in freshwater environments are maintained at migration-selection balance in the larger oceanic population and therefore available to be utilized during subsequent adaptation to new freshwater habitats. (The alleles conferring adaptation to freshwater environments are thereby “transported” to and reassembled in new lakes.) The first clear example of the global reuse of such alleles in stickleback was the gene *eda* which has been shown to be an important regulator for the number of bony lateral plates [Colosimo et al., 2004, Cresko et al., 2004, Colosimo et al., 2005]. While the low lateral plate version of this gene arose millions of years ago, it is found in much younger freshwater ponds around the Northern Hemisphere [O’Brien et al., 2015]. More recently, genome-wide haplotype analyses have provided evidence that *most* regions of the genome that distinguish marine-freshwater genetic differences share this pattern [Nelson and Cresko, 2017].

While the growing body of population genomic data on stickleback evolution supports the transporter hypothesis, a number of questions remain. What are the actual population sizes, migration rates, and fitness differentials consistent with this hypothesis? How many differentially selected alleles exist, how many are used at any one time, and how are they arranged within the genome? A curious natural history observation underlying many of these questions is the fact that some newly formed freshwater habitats, such as the ponds on Middleton Island, are quite small and presumably the number of initial marine migrants is few. The variation carried by these few initial migrants might therefore be a small subset of the total variation, and thus be insufficient to fuel adaptation without subsequent influx of alleles from additional generations of marine migrants carrying the remaining freshwater adaptive alleles.

Here, we use individual-based forward simulations implemented in SLiM that incorporates selection on a quantitative trait explicitly determined additively from a realistically long genome to model the stickleback metapopulation and address these questions [Haller and Messer, 2017, 2018]. We ask how variation in amount of gene flow affects the genetic architecture of local adaptation to newly created freshwater ponds. Because we record the entire genealogy of all alleles [Kelleher et al., 2018], we can determine the distribution and abundance of haplotypes across all marine and freshwater populations to know the timing and proportion of potentially adaptive alleles that are actually utilized in each population. In addition, we can document how these adaptive alleles are distributed across the genomes, and as a consequence determine how this may affect the efficacy of genome scans for between-habitat differentiation.

## Methods

To explore these questions, we used SLiM [Haller and Messer, 2017, 2018] to implement forward-time simulation of populations of individuals with explicitly represented genomes in which selection acted upon a single continuous quantitative trait. The details of the model were motivated by current understanding of threespine stickleback evolutionary history and demography, and included divergent selection in the two habitats which had substantial spatial structure. Certain aspects of the model remain simplistic due to computational constraints; in particular, total population sizes are much smaller than in reality, although we may capture a good picture of local dynamics.

### Habitat and geography

Our simulations include two habitat types – marine and freshwater – defined by the nature of their selective pressures, each with 5,000 diploid individuals. The arrangement of these habitats, depicted in Figure 1, roughly models a set of freshwater habitats along a stretch of coastline. The marine habitat is a continuous, one-dimensional range of length 25 units, while the freshwater habitat is divided into 25 discrete subpopulations (which we call “lakes”), each connected to the marine habitat at regularly spaced intervals (positions *i* − 1*/*2 for 1 ≤ *i* ≤ 25).

**Figure 1:**
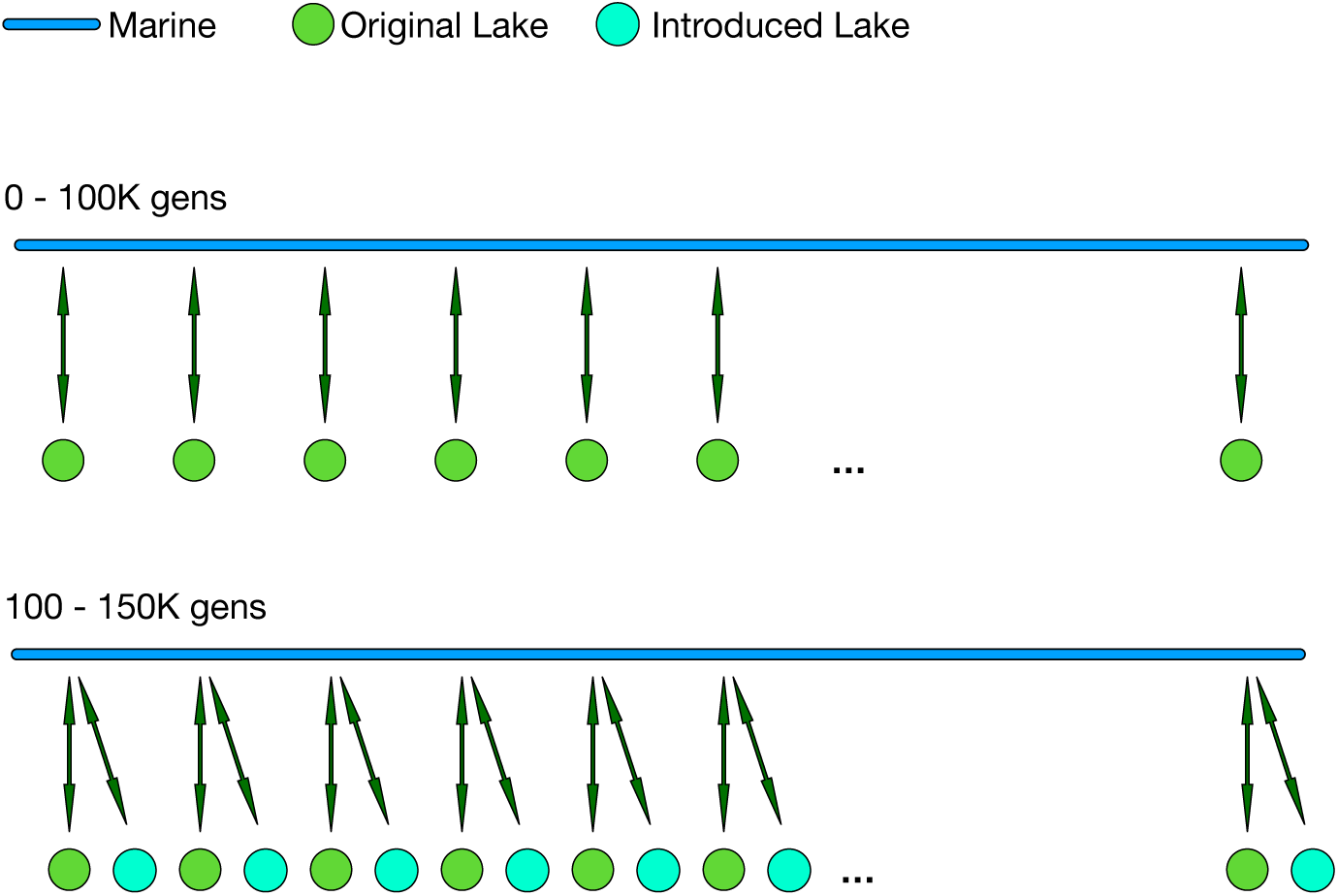
**Diagram of simulated populations:** a single, continuous, one-dimensional marine habitat (blue) is coupled to “lakes”, in which stickleback randomly mate, at discrete locations with arrows representing migration patterns. After an initial period of 100K generations with 25 lakes, an additional 25 lakes are added (at the same set of locations) and populated with marine individuals to simulate the appearance of newly accessible freshwater habitats colonized by marine stickleback. The marine habitat, and each set of 25 lakes, each contain 5,000 individuals at all times.

Divergent selection is mediated by a single quantitative trait with different optima in marine and fresh-water habitats. This situation roughly models the cumulative effect of the various phenotypes thought to be under divergent selection between the habitats, such as armor morphology, body size, craniofacial variation and opercle shape. The optimal trait values in the marine and freshwater habitats are +10 and −10 respectively, and fitness of a fish with trait value *x*_ind_ in a habitat with optimal value *x*_opt_ is determined by a Gaussian kernel with standard deviation 15, i.e.,

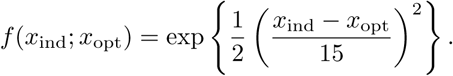

Thus, the difference between an individual’s trait value and the optimum determines that individual’s fitness. We chose the difference between optima and strength of stabilizing selection in each habitat so that (a) around 10 (diploid, homozygous) mutations were sufficient to move from one optimum to the other, and (b) well-adapted fish from one habitat would have low, but nonzero, fitness in the other habitat.

### Genetic architecture of the trait

Each individual carries two linear chromosomes, each of size 10^8^ loci. Mutations that can affect the trait under selection can occur at rate 10^−10^ per locus per generation in ten regions of 10^5^ loci each, spread evenly along the chromosome. Each mutation in these regions is either additive, completely recessive, or completely dominant (with equal probability). Effect sizes for these mutations are chosen randomly from an exponential distribution with mean 1*/*2, either positive or negative with equal probability. Individual trait values (*x*_ind_) are determined additively from the diploid genotypes. Concretely, an individual that is heterozygous and homozygous for mutations at sets of loci *H* and *D* respectively has trait value 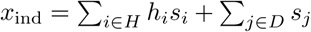, where *h*_*i*_ and *s*_*i*_ are the dominance coefficient and the effect size of the mutation at locus *i*. Subsequent mutations at the same locus replace the previous allele.

### Population dynamics

We use SLiM to simulate a Wright–Fisher population with non-overlapping generations and a fixed population size of 5,000 diploid individuals in each habitat. Each generation, the two parents of each new offspring are chosen proportional to their fitness (unlike actual stickleback, all individuals are hermaphroditic), and the contributing genomes are produced by Poisson recombination with an average of one crossover per chromosome per generation (10^−8^ per locus per generation). Since the total population across *all* 25 lakes is fixed at 5000, and the Wright–Fisher model assumes unrealistic global population regulation, we normalize the fitnesses of each individual so that approximately 200 offspring are generated in each lake, each generation. (A better implementation would use SLiM’s “non-Wright-Fisher” model type, which was not available when we wrote the simulations; but we do not expect the difference to affect results.) To do this, we divide fitness values of each freshwater individual by the mean fitness in their lake, so that the mean fitnesses of all lakes are equal before selection happens.

As depicted in Figure 1, dispersal occurs both locally along the coastline in the marine habitat as well as between the marine habitat and the lakes, with a lake–ocean migration rate denoted *m*. No migration occurs among the freshwater populations. All individual dispersal events can be thought of as occurring at the juvenile stage in the life cycle of the simulation. Each new individual in each habitat has parents from the other habitat with probability *m* (in which case we call it a “migrant”), and parents from the same habitat with probability 1 − *m*. The first parent of each non-migrant individual in the freshwater habitat is chosen from the freshwater habitat proportional to fitness, and a mate is chosen from the same lake as the first, also proportional to fitness. The offspring then lives in the same lake as the parents. Parents for each non-migrant marine individual are chosen similarly: first, a single parent is chosen proportionally to fitness in the marine habitat, and then a mate is chosen, also proportionally to fitness but re-weighted by a Gaussian function of the distance separating the two, with standard deviation 1*/*2. Concretely, if the first parent is marine individual *i*, then marine individual *j* is chosen as the mate with probability proportional to 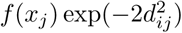, where *f* (*x*_*j*_) is the fitness of individual *j* and *d*_*ij*_ is the distance between the two locations. Finally, each new marine offspring is given a position displaced from the first parent’s position by a random Gaussian distance with mean 0 and standard deviation 0.5, and reflected to stay within the habitat range. Parents for each freshwater migrant are chosen in the same way as for non-migrant marine individuals, and are assigned to the lake nearest to the position of the first marine parent. Similarly, parents for each marine migrant are both chosen from the same lake as before, and the offspring is given a spatial location in the marine habitat at the location of the parent’s lake.

### Colonization of newly formed lakes

To study how marine-derived populations adapt after colonizing newly appearing freshwater habitats, we introduce a new set of 25 lakes after 100,000 generations. These new lakes are populated with marine individuals to emulate a freshwater lake being colonized by oceanic stickleback that had previously been exchanging alleles with older freshwater populations. This creates two *sets* of lakes, which along with the marine population have a total of 15,000 individuals. Since this introduction of lakes doubles the number of lake-to-marine immigrants, the probability that a new marine individual has freshwater parents is 2*m* instead of *m*.

### Recording genealogical history

We used SLiM’s ability to record *tree sequences* [Haller et al., 2018, Kelleher et al., 2016] to output the genealogical history of all individuals at the time of introduction of new lakes, at the time of adaptation, and at the end of the simulation. This allowed us to directly query the true origins of adaptive alleles. In addition, it allowed for much larger simulations by avoiding the computationally expensive task of simulating neutral mutations which were retroactively added to the gene trees at a rate of 10^−8^ per locus per generation, as described in Kelleher et al. [2018]. The tree sequence output by each simulation allows us to explore the origin of the genetic basis of adaptation in the new lakes. To do this, we constructed the genealogical tree relating all extant chromosomes at each locus along the genome. Using these trees we classified each adaptive allele, in each genome in the new lakes at the time of adaptation, into four categories:

1. *a* “*De novo*” *allele:* deriving from a new mutation that occurred in a new lake.
2. *a* “*Migrant*” *allele:* deriving from a migrant not in the initial generation that colonized the lake
3. *a* “*Captured*” *allele:* present in initial colonists of the new lake, and both common (above 50%) in the original lakes, and uncommon (below 50%) in the ocean.
4. *a* “*Marine*” *allele:* present in initial colonists of the new lake, but not a “captured” allele.

The proportion of trait-affecting alleles in new lakes that fall in these categories measures the degree to which selection in the new environments made use of (1) new mutation, (2) post-colonization migration, (3) standing variation at migration–selection balance, and (4) standing variation at mutation–selection balance.

We used neutral mutations to calculate *F*_*ST*_ for each locus, which we then averaged in windows as a measure of between-population relative differentiation. Concretely, if *p*_*f*_ and *p*_*m*_ are the frequencies of a given mutant allele in the freshwater and marine habitats, respectively, and 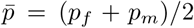, then we compute *F*_*ST*_ for that mutation as 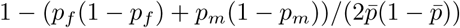.

## Data availability

The source code used to run and analyze the simulations in this paper are available at https://github.com/jgallowa07/SticklebackPaper.

## Results & Discussion

To observe the impact of gene flow on selection in the new lakes, we varied the ocean–lake migration rate, *m*, across separate simulations from 5 × 10^−5^ to 5 × 10^−1^. Below, we often refer to these migration rates in terms of the number of migrants per lake, per generation, which we denote *M*. Since each lake contains 200 individuals, *M* = 200*m*. Many aspects of adaptation changed substantially across this range, including the speed of adaptation, degree of sharing of adaptive alleles between lakes, and the population genetic signals left behind. At very low rates of gene flow, each new lake’s population adapted almost completely independently through *de novo* mutation, which took a very long time (≈ 20, 000 generations). At very high rates of gene flow, local adaptation was constrained by the large influx of locally maladaptive alleles. Between these two extremes, genetic variation that allowed adaptation to freshwater habitats could move relatively easily between lakes. Perhaps surprisingly, only a few migrants per generation from the lakes to the ocean were needed to maintain sufficient genetic variation in the ocean to dramatically accelerate adaptation in new lakes.

### Rapid local adaptation at intermediate gene flow rates

Local adaptation occurred in all simulations with the exception of the highest gene flow at which half of each population was composed of migrants, as can be seen by the mean trait values of Figure 2. Figures 3 and S3 show how mean trait values in freshwater and marine populations diverged over time, until the trait means were close to the optimal values in each habitat. Establishment of new alleles in the lakes during the first few thousand generations manifests as jumps in the mean trait value which move the trait by an amount of order 1 every few hundred generations (Fig. 3). At the lowest rate of gene flow, *M* = 0.01, differences at around 20 commonly polymorphic sites (about 10 that shift the trait in each direction) were responsible for most of the adaptive differences between freshwater and marine habitats. As expected, increasing migration rate decreased differentiation between habitats: as seen in Figure 4, *F*_*ST*_ between marine and freshwater habitats at neutral sites steadily declines as migration increases.

**Figure 2:**
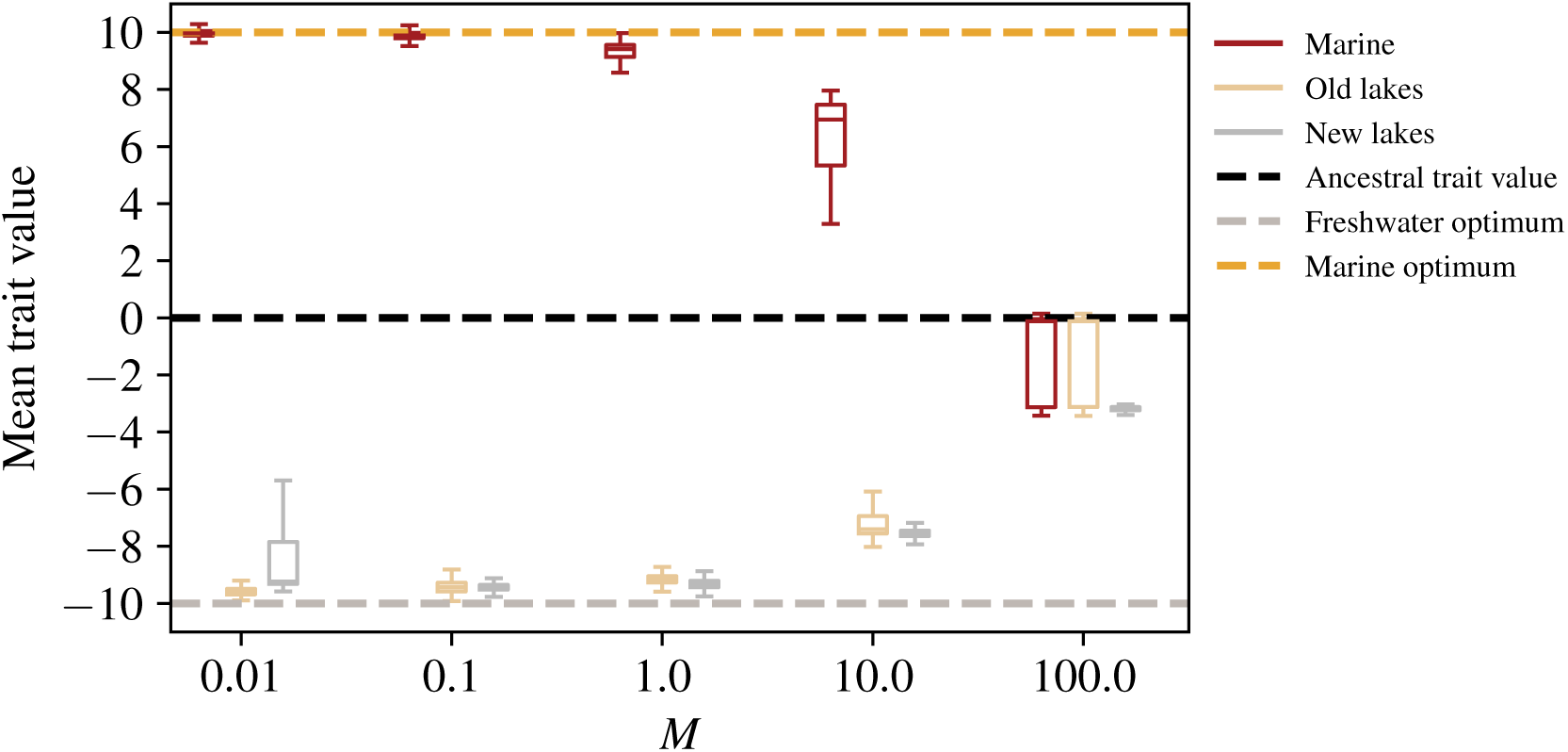
Distribution of mean individual trait values across generations of the simulation, for different migration rates. The dashed pink and purple lines at ±10 give the optimum phenotypes in the marine and freshwater environments, respectively.

**Figure 3:**
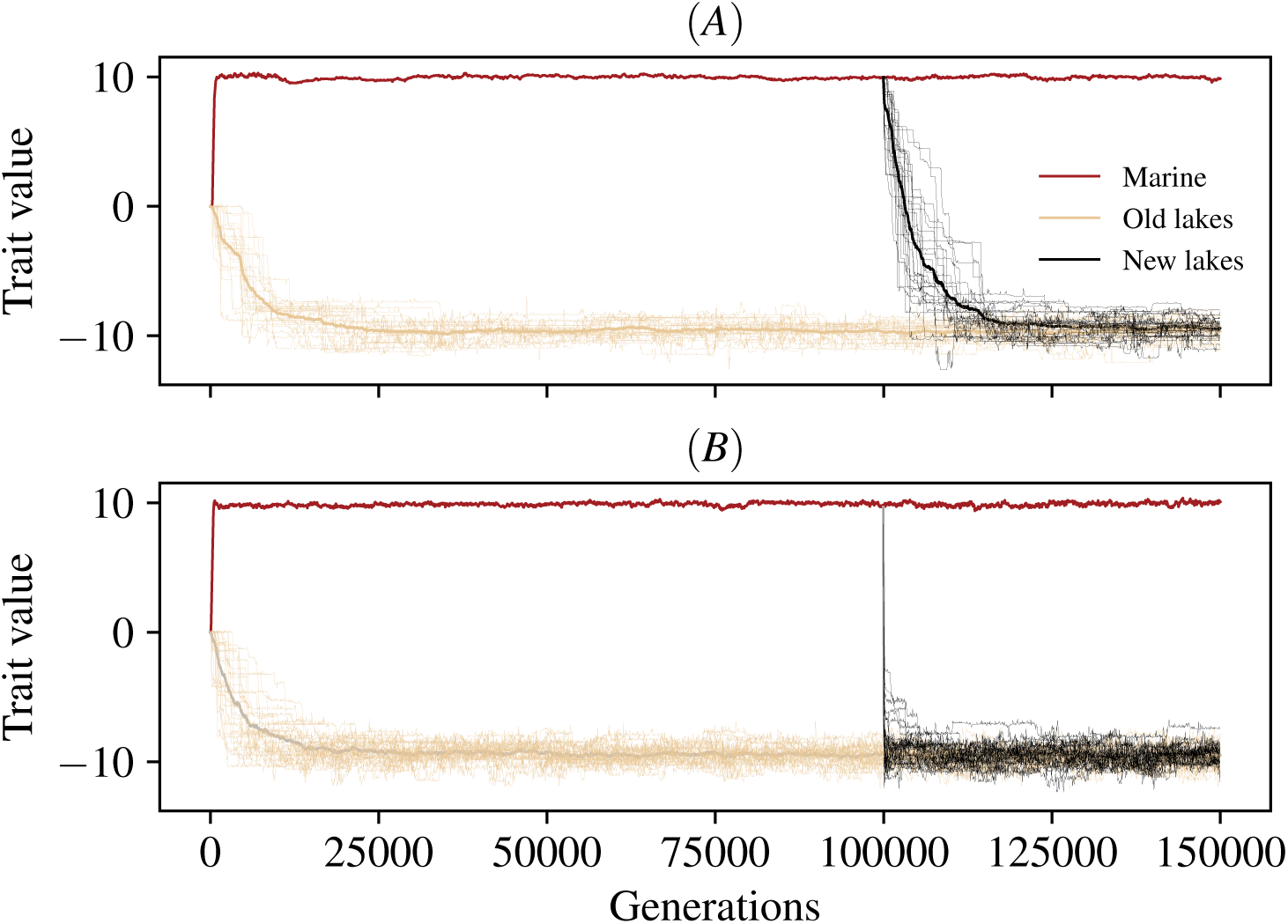
Mean individual trait values in the marine habitat (red line), the original lakes (tan lines; average thicker), and the new lakes (black lines; average thicker), across the course of two simulations, with migration rates of **(A)** *M* = 0.01 and **(B)** *M* = 0.1 migrants per lake per generation, respectively. Optimal trait values in the two habitats are at ±10. Analogous plots for other migration rates are shown in Figure S3.

**Figure 4:**
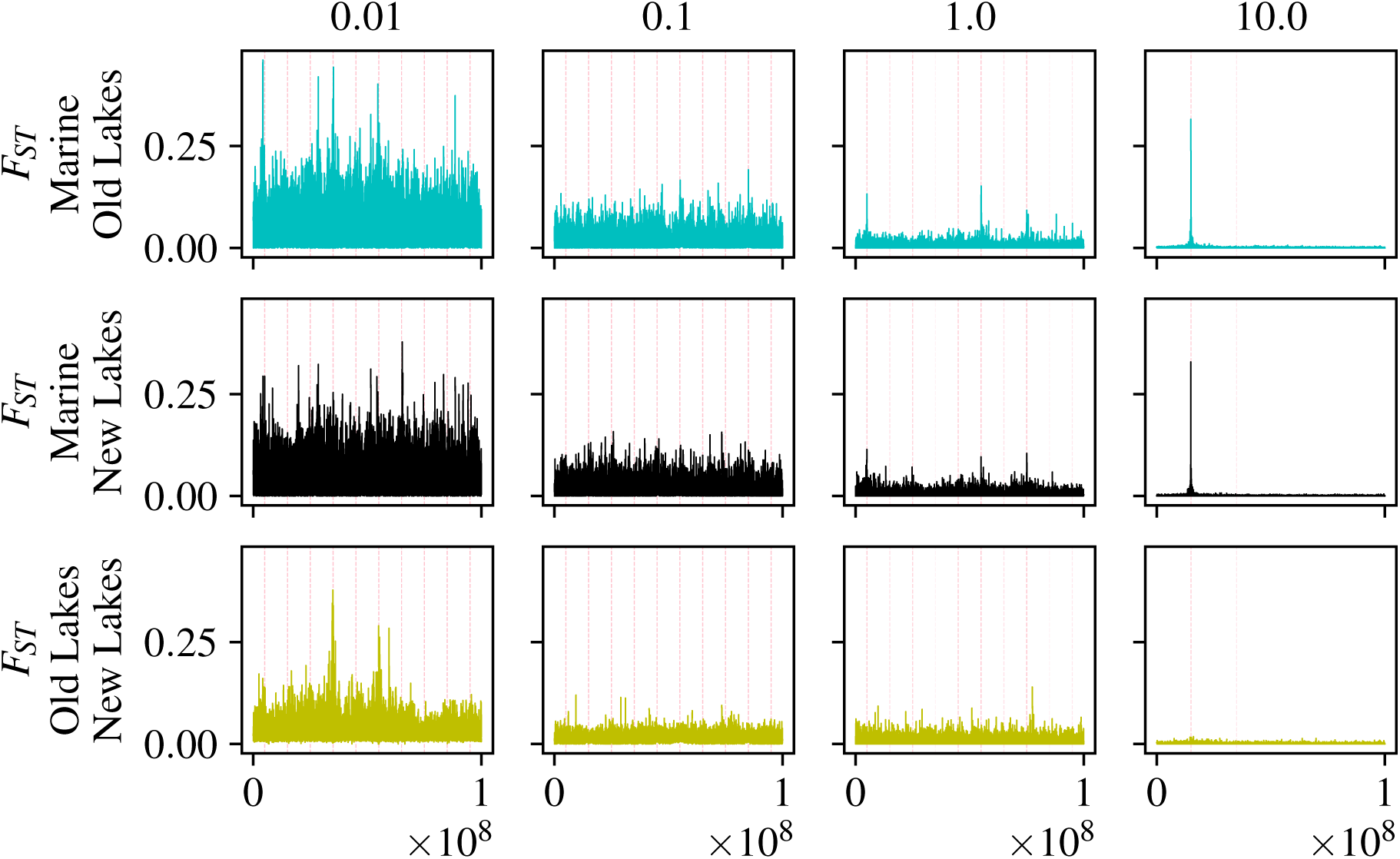
Average *F*_*ST*_ in windows of 500Mb between: **(top)** marine habitat and old lakes; **(middle)** marine habitat and new lakes; and **(bottom)** old and new lakes. Each plot shows *F*_*ST*_ values for a separate simulation, with columns corresponding to increasing gene flow from left to right. *F*_*ST*_ is calculated between all marine individuals, and all lakes pooled together. All locations of pre-existing freshwater adapted alleles have been highlighted by partially transparent pink dashed vertical lines, so darker shades of pink show more freshwater adapted alleles at that location.

Adaptation occurred much more quickly at higher migration rates, both in the old and new sets of lakes. We measured this “time to adaptation” as the number of generations until average trait values in old and new lakes were within 0.5 of each other, shown in Figure 5 for different rates of gene flow. Adaptation of new lakes took over 18,000 generations at the lowest rate of *M* = 0.01, while at *M* = 1 migrant per lake per generation, new lakes adapted in just under 60 generations.

**Figure 5:**
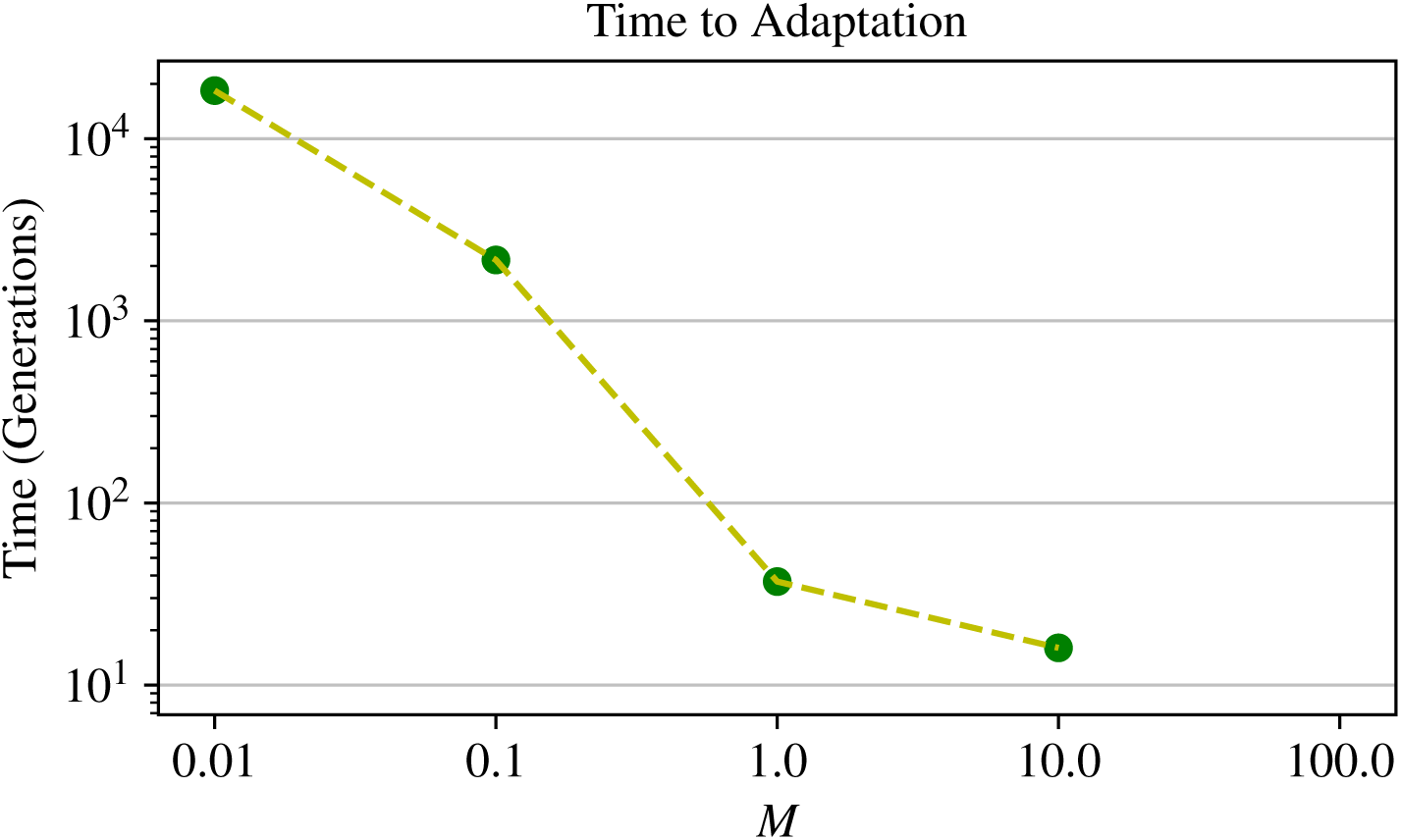
Time to adaptation as a function of migration rate. The time to adaptation is measured as the number of generations until the introduced population’s mean phenotype comes within 0.5 of the original lakes average phenotype. Each point represents a single simulation run. (Adaptation did not occur at the highest rate of gene flow.)

### Widespread allele sharing between lakes

Adaptive alleles were shared between lakes at many rates of gene flow, but not at the lowest. At low migration rates, the *initial* period of adaptation takes roughly 25 times longer for lakes than it does for the ocean. This difference occurs because in the absence of gene flow, each lake must wait for its own novel mutations to arise in order to adapt. Because the marine habitat is continuous, with 25 times more individuals than any one lake, there is a much larger influx of new mutations to be selected upon. At higher migration rates, greater mixing allows the initial lakes to share alleles instead of developing their own genetic basis for adaptation.

To investigate in more depth how locally adaptive alleles found in the original lakes are shared among newly derived lakes, as well as how they spread to the new lakes, we defined and tracked the distribution of “pre-existing freshwater adapted alleles” at the beginning of each generation. To be considered in this category, an allele must participate in the genetic basis of local adaptation for at least one of the original lakes. Concretely, these are any non-neutral mutations whose frequency is above 50% in at least one original lake and below 50% in the marine habitat. (So, “captured” alleles as defined above are pre-existing, but not all pre-existing alleles are captured.) Figure 6A shows the distribution of the number of these alleles across generations. At *M* = 0.01, each lake has a private set of about 10 mutations nearly fixed in that lake but not elsewhere: new lakes independently acquire new adaptive alleles rather than pre-existing ones. At *M* = 0.1, we again observe the original lakes adapting nearly independently from each other, but now the new lakes adapt using pre-existing alleles present in the original set of lakes. Concurrently, the average marine individual carries ≈ 2 pre-existing freshwater adapted alleles, standing variation which was nearly absent at *M* = 0.01. As migration rate increases past this, the total number of pre-existing freshwater adapted alleles declines (Figure 6C). Interestingly, the frequency of these alleles in the ocean stays relatively constant across the reasonable rates of gene flow.

**Figure 6:**
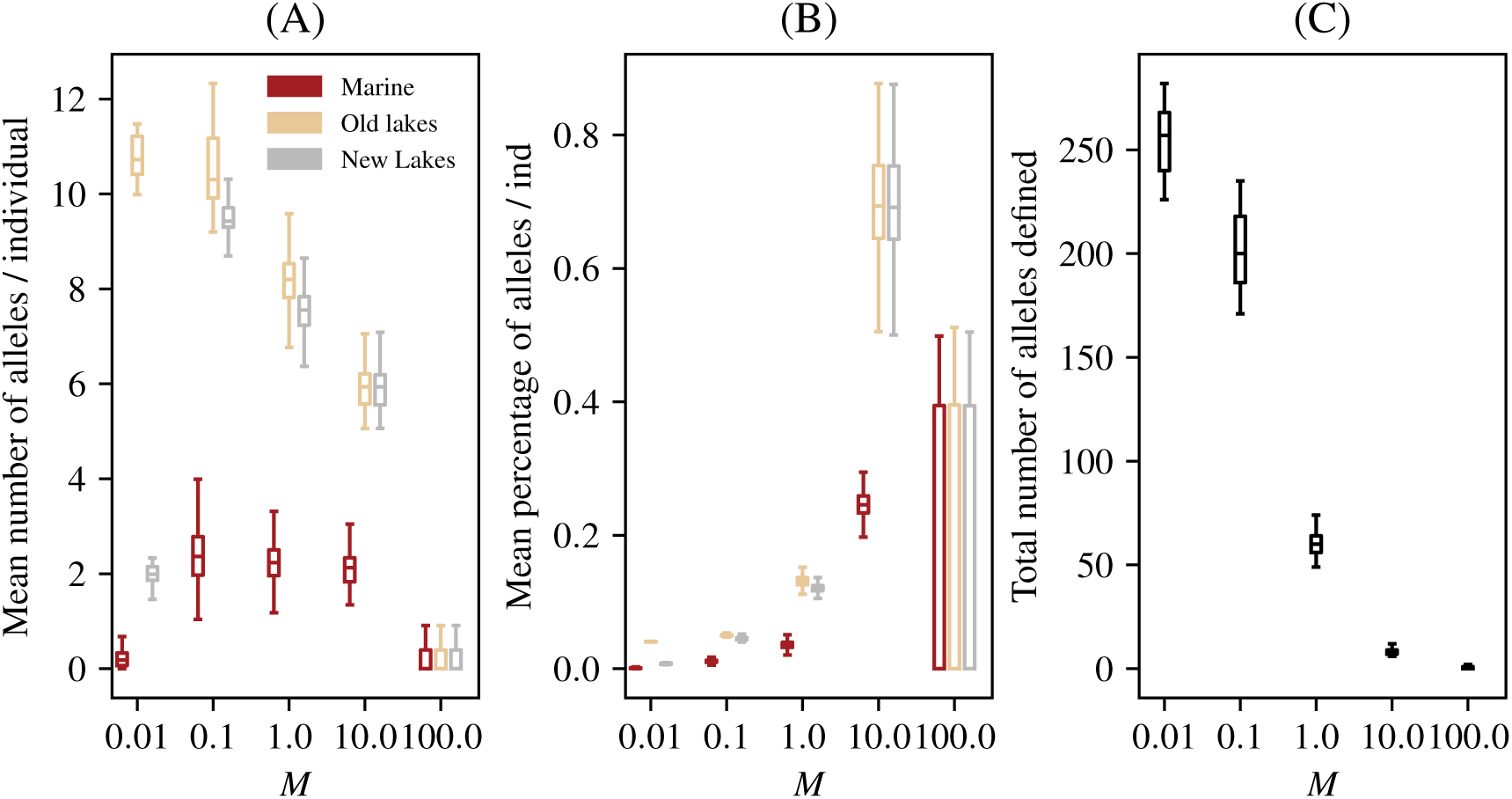
Amount of standing freshwater variation by habitat, across migration rates. Each plot counts “pre-existing freshwater adapted alleles”, that are common in the original lakes but rare in the ocean (see text for definition). **(A)** Mean number of these alleles per individual. **(B)** Mean percentage of these alleles per individual. **(C)** Total number of these alleles (so, *B* = *A/C*). The number of alleles meeting these conditions changes over the course of the simulation, and each plot shows distributions of these values throughout each simulations. The horizontal axis shows *M*, the mean number of migrants per lake per generation.

Figure 6B shows the distribution, through time, of the mean *percentage* of currently-defined freshwater adapted alleles that each genome in each of the populations carries. If all individuals across lakes carried the same set of alleles determining their trait value, this would be 100%. At the lowest migration rate (*M* = 0.01), each genome in the original lakes have almost exactly 1*/*25^*th*^ of the total number of pre-existing freshwater adapted alleles – this is because each one of the 25 lakes has adapted with a unique set of alleles. Since these are *pre-existing* alleles, the value is zero for introduced lakes. Figure 6A shows us that at 0.1 migrants per lake per generation and above, the average individual across the new lakes has nearly the same amount of pre-existing freshwater adapted alleles as individuals across the old lakes. As expected, the genetic basis of the freshwater phenotype seems to simplify as migration increases – higher rates of migration allow adaptive alleles of larger effect to travel more efficiently through the population, even though they are deleterious in the ocean.

The numbers in Figure 6 strongly suggest that the dramatic increase in speed of local adaptation we observed above occurs because higher gene flow allows sharing of freshwater alleles among populations. We confirmed this by using recorded tree sequences to identify the origin of each trait-affecting allele in each individual in the new lakes, as defined in the Methods. Figure 7 shows that at the lowest rate of gene flow the majority of adaptive alleles are derived from “*de novo*” mutation. As gene flow increases, a larger fraction of adaptive alleles derive from pre-existing variation in the marine population at the time of introduction. In other words, greater mixing at higher migration rates allows lakes to share alleles instead of developing their own genetic basis of adaptation.

**Figure 7:**
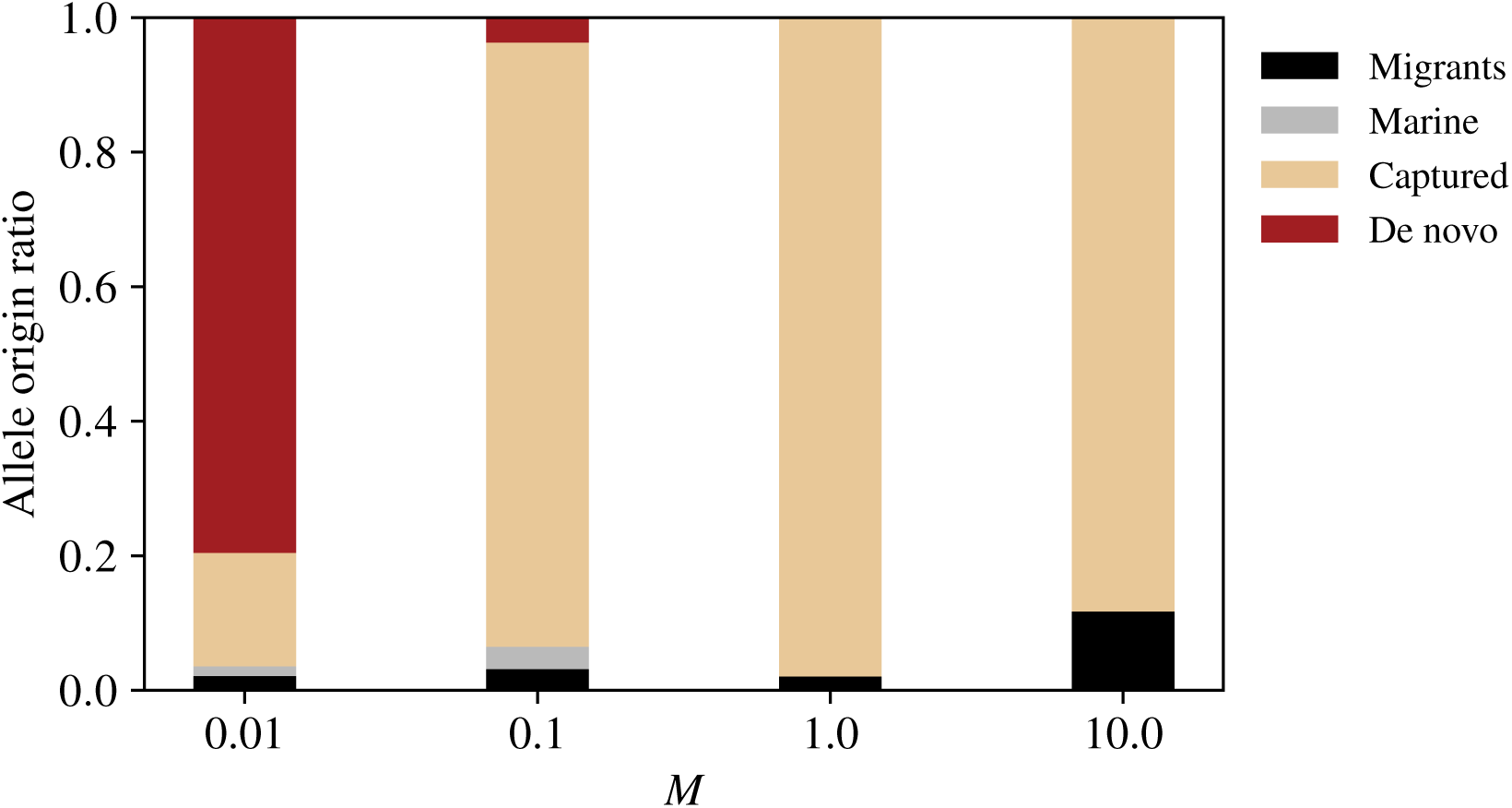
**(Origin of adaptive alleles:)** Each bar plot shows the origins of all trait-affecting alleles above frequency 50% in at least one new lake, classified as **(red)** new mutations, **(blue)** post-colonization migrants, **(green)** “captured” from pre-existing lakes, or **(orange)** standing marine variation. See Methods for precise definitions of these categories.

While increased migration allows sharing of adaptive alleles among lakes, at *M* = 10.0 migrants per lake per generation the constant influx of alleles between the habitats creates substantial migration load. This level of gene flow only replaces 5% of each population each generation with migrants from the other habitat, but is sufficient to shift the mean trait values to nearly half their optimal values, as seen in Figure 2.

### The efficacy of genomic scans of selection depends on gene flow

Here, we take a closer look at the genomic architecture of local adaptation between the two habitats. Can measures of local differentiation such as *F*_*ST*_ be used to identify the causal loci? Figure 4 shows plots along the genome of average per-locus *F*_*ST*_ values in 500bp windows between the marine habitat and all freshwater habitats pooled together. Higher rates of dispersal showed more distinct *F*_*ST*_ peaks over polymorphic loci, while “background” levels of *F*_*ST*_ increase as gene flow decreases, swamping out this signal until the regions under selection are indistinguishable. This is likely the contribution of two separate forces of natural selection: first, stronger genetic drift with less migration leads to higher background *F*_*ST*_, and second, greater sharing of adaptive alleles providing a shared signal across populations.

At first glance, this suggests that genome scans for local adaptation based purely on measures of differentiation will only be successful given enough migration between habitats. But how many of these peaks are actually underlying trait differences that form the basis for local adaptation? To quantify this, Figure S1 shows the power and true positive rates that would be obtained by an *F*_*ST*_ cutoff that declared everything above a certain value to be a causal locus.

What we observe most plainly in this graph is that *M* = 1.0 and *M* = 10.0 migrants per lake per generation are the only rates of dispersal at which a large percentage of peaks (above *F*_*ST*_ = 0.05) actually lie on top of regions which impact trait differences. Unfortunately, the low statistical power for all rates of *M* would seem to suggest that many regions which impact trait differences do not appear as *F*_*ST*_ peaks. In other words, with high rates of dispersal between population, *F*_*ST*_ peaks may reliably identify causal loci, but not all causal loci will appear as *F*_*ST*_ peaks. It’s important to note that *F*_*ST*_ in Figures 4 and S1 was calculated after pooling together all lakes and all marine individuals. For comparison, Figures S4 & S2 shows *F*_*ST*_ calculated with individuals only in the first fifth of the range (5 lakes and corresponding marine habitat). These show a substantially lower true positive rate, as population-specific noise swamps any signal between the lakes.

As has been suggested elsewhere, introgression aids identification of locally adaptive loci from scans for differentiation. This is generally because shared adaptive alleles provide something akin to independent replicates, allowing signal to emerge above the noise created by genetic drift. However, the required sample sizes and conditions on levels of admixture may be restrictive in practice. More generally, the difficulty of correctly identifying causal loci in even our most ideal situation calls for caution (and proof-of-concept simulations) in designing and carrying out scans for alleles underlying locally adaptive, polygenic traits.

### Alleles for freshwater adaptation are mostly present in the initial generation

We have thus far found that the speed of adaptation depends strongly on the degree to which alleles can be shared between populations. However, the *origin* of the alleles underlying the phenotype is still unknown. What is the nature of selection on standing genetic variants? In our simulations migration from marine individuals into the new lakes continues throughout introduction and adaptation: but is consistent influx of standing genetic variation a necessity for rapid adaptation of the population? To answer this, we traced the genealogy of *all* adaptive alleles in *all* individuals from the introduced lakes, after local adaptation. Surprisingly, for all cases with the exception of *M* = 0.01, we found that the majority of adaptive allele origins traced back the original generation of inhabitants in the lake (Figure 7). In our simulations, any reasonable subset of the ancestral population has the potential for rapid adaptation without the need post-colonization hybridization events.

If effectively capturing standing genetic variants is the key to rapid adaptation, as we have presented thus far, why is there a large difference in speed of adaptation between dispersal rates of *M* = 0.1 and *M* = 1.0 seen in Figure 5? This is surprising because we see a similar quantity of allele sharing at *M* = 1.0, but much more rapid adaptation (≈ 60 versus 2000 generations).

A possible explanation is that at lower rates of gene flow, freshwater alleles present as standing variation in the marine habitat are more tightly linked to marine alleles. This implies that adaptation in the new lakes must wait for recombination to separate freshwater and marine adapted alleles. This might be expected at low dispersal rates since freshwater alleles may need to be “masked” by nearby compensatory alleles to remain in the ocean for long periods of time. Reversing this “masking” then slows the process of adaptation that uses this genetic variation.

In other words, higher dispersal from freshwater to the ocean maintains relatively intact freshwater haplotypes that can be more easily rebuilt in the marine environments. Recall that trait-affecting mutations only occur in relatively small regions of 10^5^ loci in which recombination occurs only once in every one thousand meioses. This implies that even if there is a sufficient amount of variation in the initial population of a lake to shift the trait from the marine optimum (+10) to the freshwater optimum (−10), rebuilding the most beneficial haplotype may prove to be a non-trivial task for selection. For example, suppose there are 10 variants segregating at low frequency with effect size −1 each, but each is paired with a compensatory allele with effect size +1. Each local haplotype is therefore neutral. This might also explain why at *M* = 0.1, marine individuals still hold on average several alleles that shift the trait in the freshwater direction (Figure 6A).

To quantify the genetic variation available *without* recombination within the ten genomic regions, we first found, within each population, the haplotype with the largest net negative effect at each of the ten genomic regions. Summing these ten numbers, we get the maximum amount that selection could move the population in the freshwater direction without recombining within these regions. The mean of this value across the 25 lake populations is shown in Table 1 – typical populations at *M* = 0.1 migrants per lake per generation could shift to a phenotype of −12.02 (and so have sufficient variation to adapt without recombination), but the “best” haplotypes in the populations at *M* = 1.0 have effect sizes nearly twice as big at a mean total of −21.02. The amount of variation available at *M* = 10.0 is lower (only −7.02), presumably because of migration load, while at *M* = 0.01 almost no alleles with negative effect are present. Note that some of these haplotypes will likely be lost to drift – indeed, if they did all fix, then populations at *M* = 0.1 would adapt much more quickly. However, this calculation supports our explanation above: it appears that adaptation at *M* = 0.1 is slower because it must wait for intralocus recombination to free up genetic variation present but masked in the marine population.

**Table 1:**
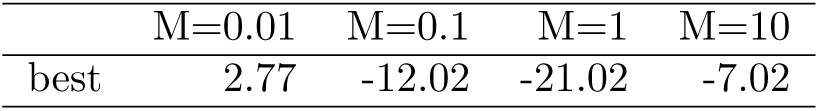
Haplotypic variation present in the new lakes at time of colonization, across rates of gene flow, **“Best”** quantified as the most negative trait value achievable with intact haplotypes, averaged across populations (see text for details).

### Simulation results align with theoretical expectations

How do our results compare to what is expected from population genetics theory? Our simulations included many loci under selection and in linkage to many other selected alleles, which makes precise calculation impossible. Nonetheless, rough calculations based on simple population genetics theory – with the benefit of hindsight – turn out to describe qualitatively most of the aspects of adaptation we observed above. Because we model stabilizing selection on an additive trait controlled by a moderately large number of loci within each population, more precise expectations might be obtained through quantitative genetics [Svardal et al., 2014] or even Fisher’s geometric model [Barton, 2001, Chevin et al., 2014], but doing so is beyond the scope of this paper

One of the most useful things that theory tells us is about the fate of a new allele in a lake, that has appeared by either migration or mutation. If the allele has fitness advantage *s* – – i.e., when it is rare but present in *n* copies, the expected number of copies in the next generation is (1 + *s*)*n* – then the probability that it escapes demographic stochasticity to become common in the population is approximately 2*s* [Lambert, 2006, Haldane, 1927], assuming Poisson reproduction, as we roughly have here. Since we are studying a quantitative trait, the fitness effect of each allele depends on the population context: if the individuals in the population all have trait values *z* units above the optimum, and the allele has effect size −*u* in heterozygotes, then the fitness advantage of the allele is the ratio of fitnesses with and without the allele. As we calculate fitness here, this is *s*(*u*) = exp(−*β*((*z* − *u*)^2^)*/* exp(−*βz*^2^)) ≈ 2*βzu*, where *β* = 1*/*450. This tells us two things: (1) the rate of adaptation decreases as the population approaches the optimum, and (2) larger mutations (in the right direction) are more likely to fix.

#### New mutations

The total rate of appearance of new mutations per lake is *µ*_*L*_ = 0.04 per generation, and these are divided evenly in six categories: additive, dominant, and recessive, in either direction. Therefore, a new additive or dominant effect mutation appears once every 75 generations, on average. The effect size of each new mutation is randomly drawn from an Exponential distribution with mean 1*/*2, and so averaging the probability of establishment over this distribution, we get that the probability that a dominant mutation manages to establish in a population differing from the optimum by *z* is roughly 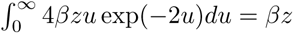. Multiplying the rate of appearance of these mutations with the probability they establish predicts that the rate of establishment of dominant mutations is *βz/*75, i.e., about one such mutation every 33750*/z* generations. During the initial phase of adaptation, the populations begin at around distance *z* = 10 from the optimum, and *z* decreases as adaptation progresses. Mutations that successfully established are more likely to be strong: the distribution of the effect sizes of these successfully established mutations has density proportional to *u* exp(−2*u*), which is a Gamma distribution with mean 1 and shape parameter 2. There are also additive alleles: these have half the effect in heterozygotes, and so roughly half the probability of establishment. Combining these facts, we expect adaptive alleles to appear through mutation within lakes at first on a time scale of 3,000 generations, with the time between local fixation of new alleles increasing as adaptation progresses, and each to move the trait by a distance of order 1. This agrees roughly with what we see in Figures 3 and S3.

#### Standing variation

How are freshwater alleles maintained in the ocean, where they are deleterious? Assuming that fish in the marine environment are close to their phenotypic optimum, an allele that when heterozygous moves the trait *u* units in the freshwater direction, has fitness roughly exp(−*βu*^2^) ≈ 1 − *βu*^2^, i.e, a fitness differential of *s* = *βu*^2^. The product of population size and fitness differential in the marine environment for a mutation with *u* = 1 is therefore 2*Ns* ≈ 22, implying that these alleles are strongly selected against but might drift to moderate frequency if recessive. The average frequency of such an allele in the marine environment at migration-selection equilibrium is equal to the proportion of individuals in the ocean replaced by migrants per generation divided by the selective disadvantage, i.e., around *m/βu*^2^. Each new lake is likely to contain a few copies of alleles at frequency above 1/400 (since each new lake is initialized with 400 randomly selected genomes). For a typical allele with effect size *u* = 1, the equilibrium migration-selection frequency is greater than this threshold if *m/β >* 1*/*400. Since 1*/β* = 450, if *m* ≥ 5 × 10^−6^ this suggests that there is a good chance that any particular lake-adapted allele that is present in all pre-existing lakes will appear at least once in the fish that colonize a new lake. However, an allele with effect size *u* = 1 only has probability of around 1/20 of establishing locally, and so must be present in about 20 copies to ensure establishment. Putting these calculations together, we expect migration-selection balance to maintain sufficient genetic variation for new lakes to adapt if *m* ≥ 10^−4^, which corresponds to *M* ≥ 0.4. This is in good agreement with our observations. However, this calculation treats each allele independently; in practice we found that standing freshwater variation in the ocean was masked by linkage to compensatory marine variants.

#### Migration

The key quantity regulating the amount of standing variation in the ocean is the *downstream* migration rate, from lakes to the ocean. How important is the upstream migration rate? If sufficient genetic variation is not present in a new lake initially, it must appear either by new mutation or by migration. Since a proportion *m* of each lake is composed of migrants each generation, it takes 1*/m* generations until the genetic variation introduced by migrants equals the amount initially present at colonization. This implies a dichotomy: either (a) migration is high, and adaptation is possible using variants present at colonization or arriving shortly thereafter, or (b) migration is low, so adaptation takes many multiples of 1*/m* generations. Since in our model lower migration also reduces the amount of variation available in the ocean, we expect very little contribution of subsequent migration across any value of *m*, as seen in Figure 7.

#### Population size

In our simulations, the marine and freshwater populations are of equal size. More generally, we expect the downstream migration rate that is required to maintain standing variation there to scale with the size of the marine population, since the same number of migrants make up a smaller proportion of a larger population. So, if the marine population was well-mixed and tenfold larger than all freshwater populations combined, the number of migrants per lake per generation would need to be ten times larger than in our simulations to maintain sufficient standing genetic variation. If the marine population was a thousandfold larger, adaptation from standing genetic variation that is deleterious in the ocean might be impossible. The fact that repeated adaptation from standing genetic variation has been shown to occur in real world threespine stickleback populations argues that the population sizes of marine and combined freshwater stickleback are not that different. While this might at first seem implausible given the ocean is large and lakes are small, in reality there are hundreds of thousands if not millions of lakes inhabited by threespine stickleback in coastal regions of the Northern Hemisphere, and the limiting habitat in the ocean constraining marine stickleback population sizes is likely to be coastal nesting areas. Together, these facts make the equivalency of marine and freshwater stickleback population sizes much more plausible.

## Conclusion

In this paper, we have analyzed a relatively realistic simulation study of a coastal meta-population to understand how repeated local adaptation operates on complex standing variation across a wide range of gene flow rates. We have shown that rapid and parallel adaptation similar to that documented in natural stickleback populations occurs over a realistic range of parameter values. These simulations therefore provide an in-depth look at (and quantitative proof-of-concept of) the “transporter” hypothesis suggested by Schluter and Conte [2009]. Selection is able to rebuild the freshwater haplotype at a rapid pace (in tens of generations) from variation present at migration-selection balance in marine populations A finding of practical consequence is that the efficacy of *F*_*ST*_-based genomic scans varies significantly with migration rate. Perhaps surprisingly, we found that the majority of adaptive variant can be traced back to the original generation of new habitats, with little role for post-colonization migrant alleles. In other words, a randomly chosen subset of marine stickleback carry the capacity to rapidly adapt in freshwater habitats without *any* continued connection to the rest of of the species. Thus, the large marine population was able to harbor and distribute alleles that were deleterious in the ocean habitat, but adaptive in freshwater habitats, to surrounding freshwater populations with only a few migrants per population, per generation.

## Supplementary material

**Figure S1:**
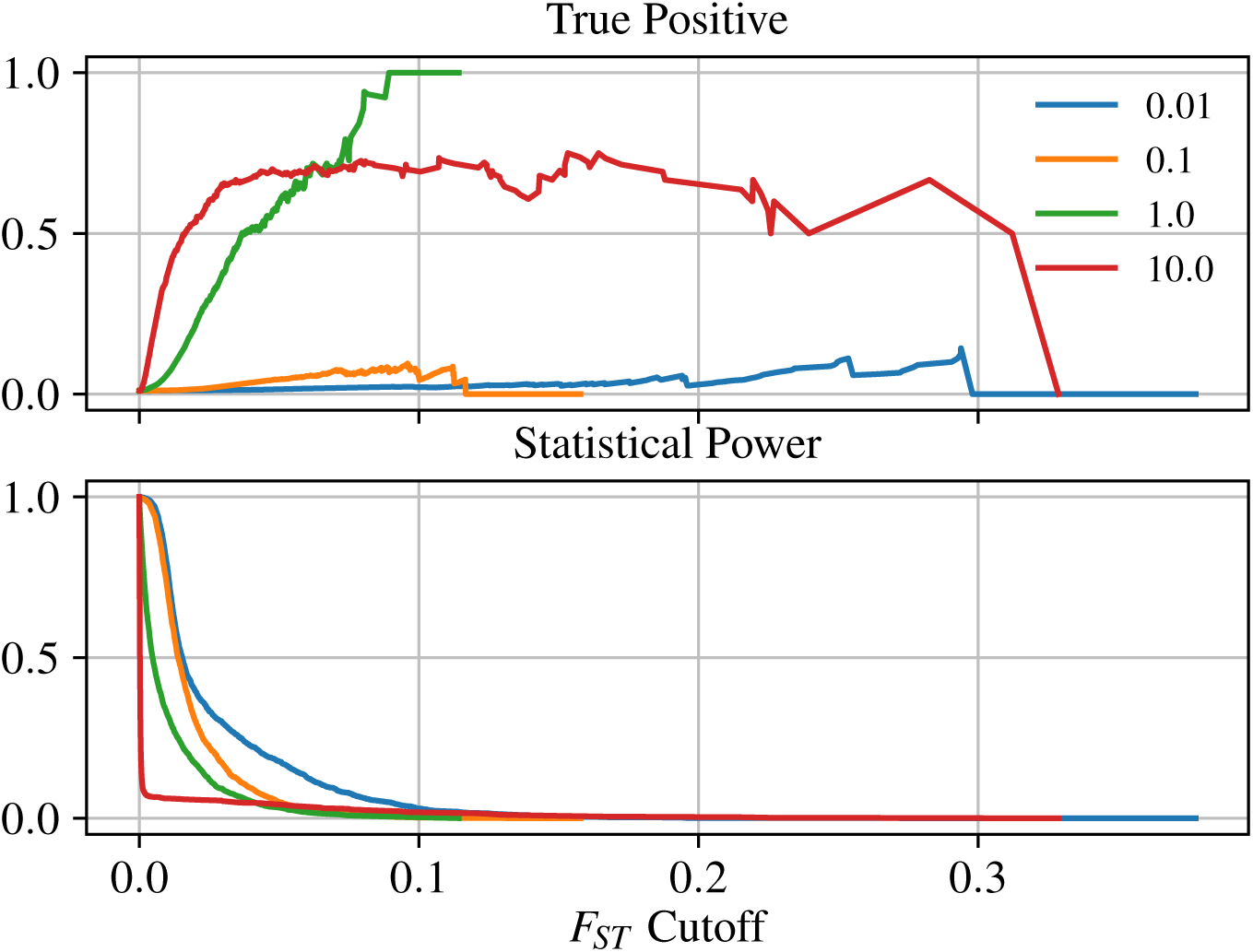
Statistical power and true positive rate as a function of *F*_*ST*_ threshold. Statistical power is the likelihood that a SNP will be predicted to have an effect on phenotype when there is an effect to be detected. True positive rate is the proportion of SNPs above the *F*_*ST*_ threshold that affect phenotype.

**Figure S2:**
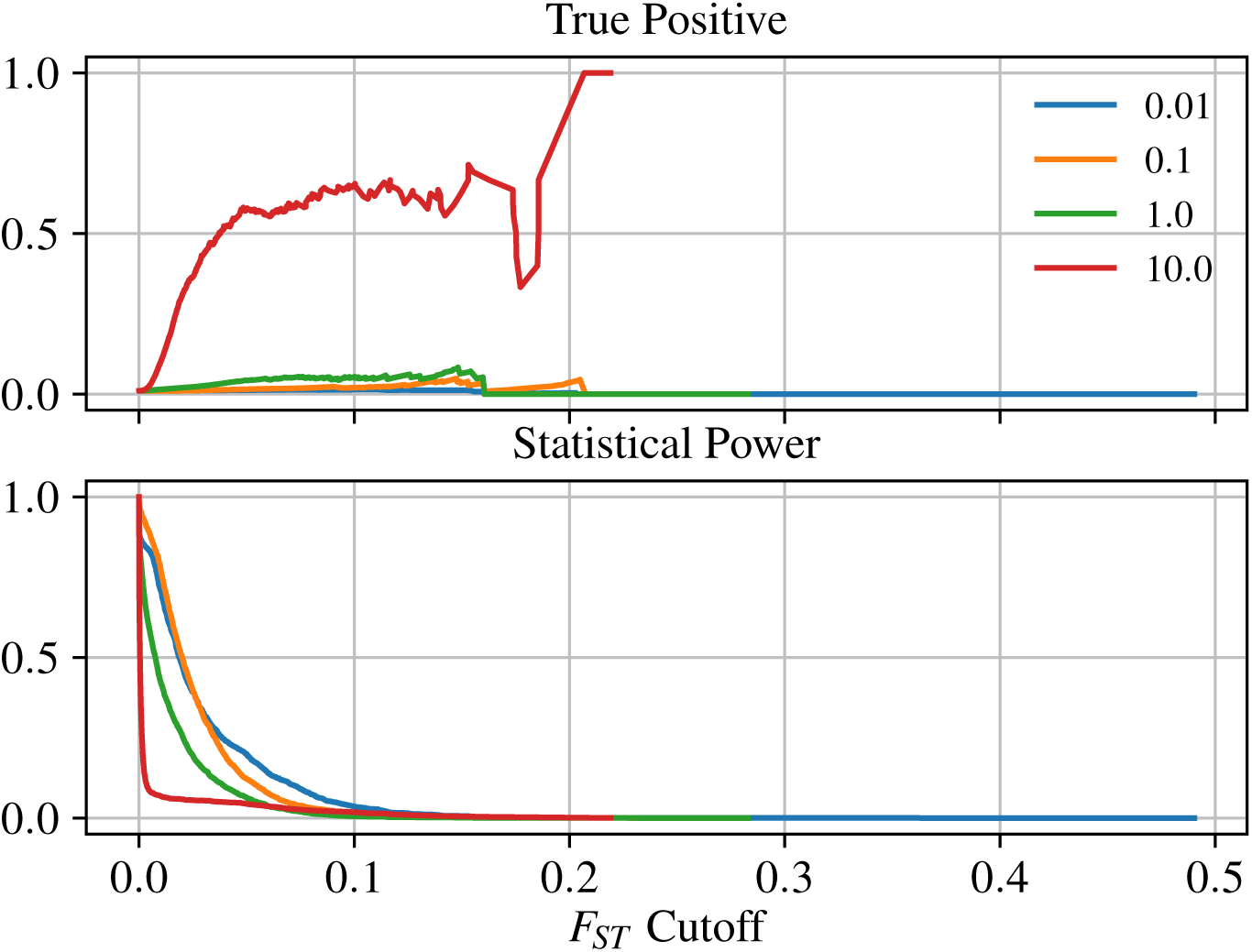
Statistical power and true positive rate as a function of *F*_*ST*_ threshold. As in Figure S1 but computed only using the first one-fifth of the total range – i.e., fish in the first five lakes and those in the ocean with coordinate less than 5.0.

**Figure S3:**
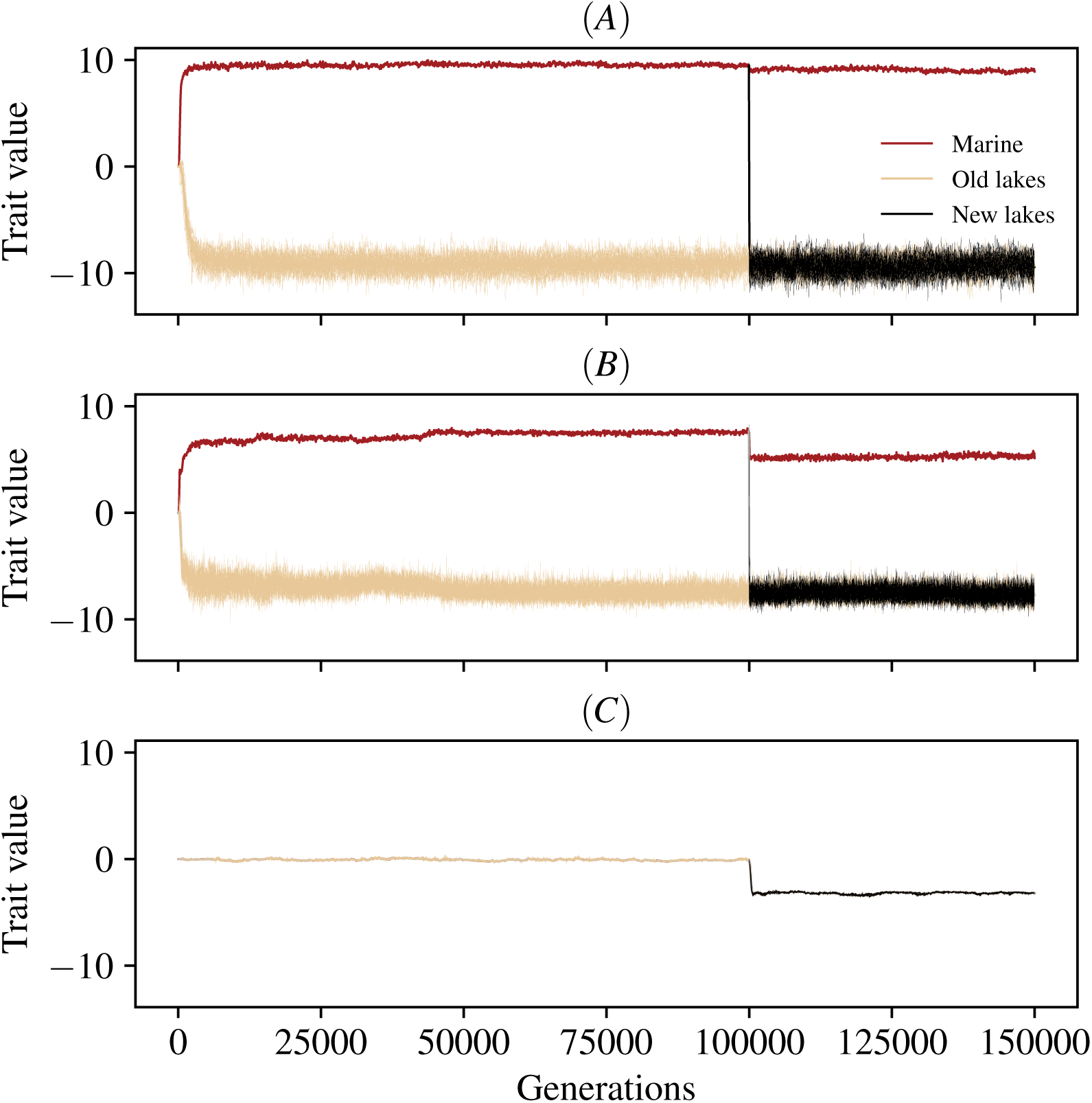
Mean individual trait values in the marine habitat (red line), the original lakes (tan lines; average thicker), and the new lakes (black lines; average thicker), across the course of two simulations, with migration rates of **(A)** *m* = 5 × 10^−3^, **(B)** *m* = 5 × 10^−2^ and **(B)** *m* = 5 × 10^−1^. (i.e., *M* = 1, 10, and 100 migrants per lake per generation, respectively). Optimal trait values in the two habitats are at ±10.

**Figure S4:**
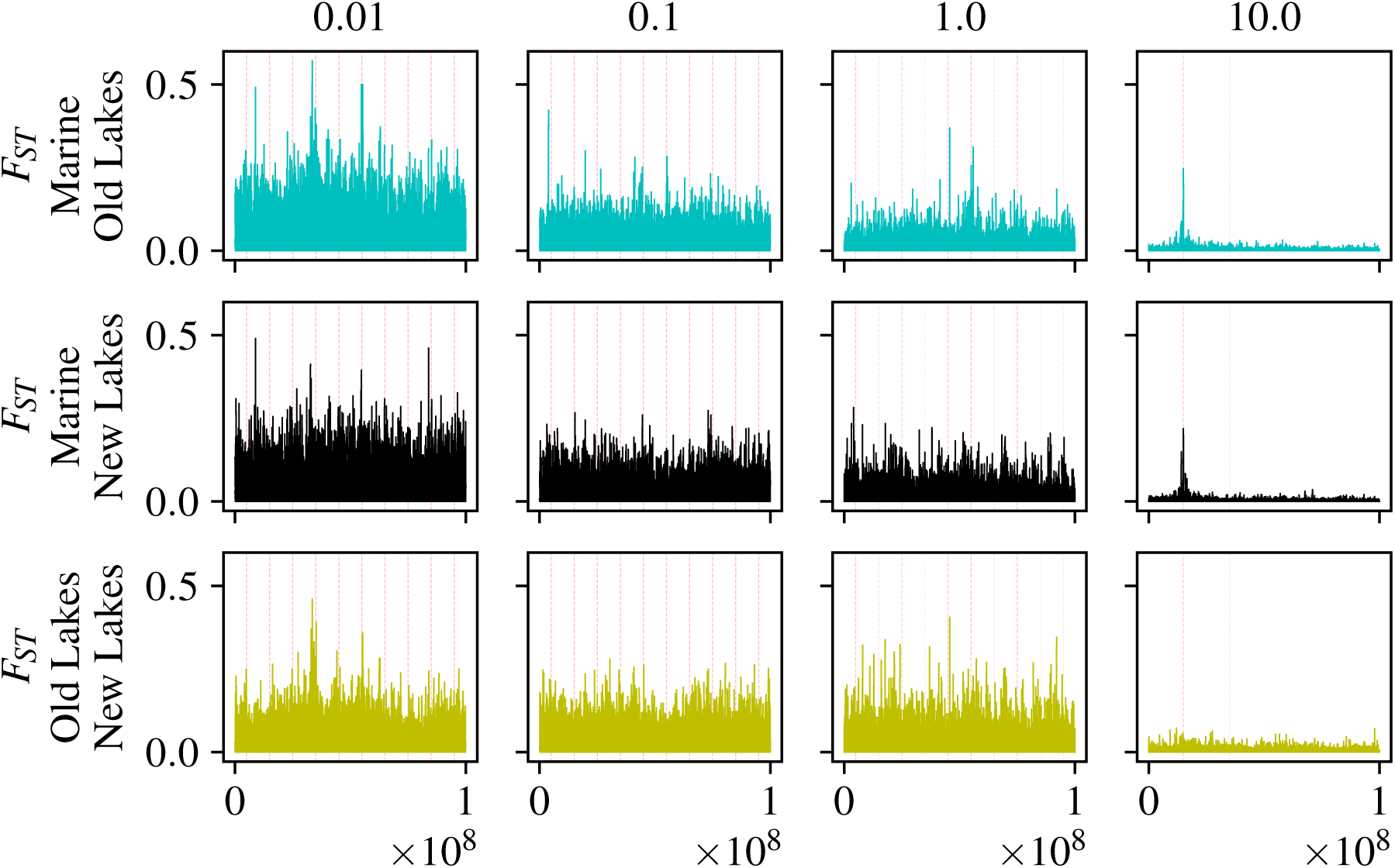
Average *F*_*ST*_ in windows of 500Mb between: **(top)** marine habitat and old lakes; **(middle)** marine habitat and new lakes; and **(bottom)** old and new lakes. Each plot shows *F*_*ST*_ values for a separate simulation, with columns corresponding to increasing gene flow from left to right. *F*_*st*_ is calculated between all marine individuals with spatial location ≤ 5.0, and the five corresponding lakes. All locations of pre-existing freshwater adapted alleles have been highlighted by partially transparent pink dashed vertical lines, so darker shades of blue imply more freshwater adapted alleles at that location.

